# Unconstrained naturalistic human brain imaging and decoding with a fully wearable high- density optical system

**DOI:** 10.64898/2026.08.24.746346

**Authors:** William T. Hamic, Wiete Fehner, Morgan Fogarty, Alvin S. Agato, Hannah E. DeVore, Sean M. Rafferty, Dana Wilhelm, Amelia M. Hines, Adam T. Eggebrecht, Jason W. Trobaugh, Edward J. Richter, Joseph P. Culver

## Abstract

Understanding how the brain supports complex cognition in real-world environments requires neuroimaging systems that impose minimal constraints on natural behavior. Existing high-fidelity modalities, such as functional magnetic resonance imaging (fMRI), confine participants to the scanner, while wearable alternatives sacrifice spatial resolution or cortical coverage. Here, we developed a fully untethered whole-head optical neuroimaging system that achieves high-fidelity tomographic reconstruction through dense spatial sampling, configurable source multiplexing, and high dynamic range detection. This wearable high-density diffuse optical tomography (WHD-DOT) system achieves 151 dB effective dynamic range and nearly 3000 source-detector measurements, comparable to the highest-performing fiber-based DOT systems, while maintaining wireless, battery-powered mobility. We validate WHD-DOT across three paradigms of increasing ecological complexity, ranging from standard functional localizers to naturalistic movie viewing and live piano performance. Across all paradigms, WHD- DOT produces robust, well-localized encoding, repeatable single-trial responses, and above-chance decoding of stimulus-specific dynamics. Piano performance, which requires continuous bimanual movement and an unconstrained posture, is a rigorous real-world test for a wearable neuroimaging system. By decoding song-segment identity of free piano performance at 71.1% accuracy (chance 12.5%), WHD-DOT shows that brain activity from unconstrained, real-world behavior, previously beyond reach of high-fidelity imaging, is now both measurable and decodable.

## Introduction

Human cognition is continuous, dynamic, and embedded in real-world contexts. The field of human neuroscience has embraced this reality by increasingly using naturalistic paradigms [1]. These range from continuous, information-rich stimuli, such as natural speech [2, 3] and movies [4–7], to interactive and social paradigms, including hyperscanning [8]. Capturing distributed cortical representations underlying these tasks requires high spatial resolution and whole-head coverage, a requirement met primarily by functional magnetic resonance imaging (fMRI). As a result, most advanced naturalistic studies have been constrained to the scanner bore. Recent advances in wearable optical neuroimaging have opened new possibilities for real-world neuroscience, allowing for unconstrained posture, natural head and body motion, and long-duration recordings in everyday environments [9–12].

Capturing distributed cortical dynamics in a wearable system remains challenging, as systems face tradeoffs among spatial resolution, head coverage, imaging fidelity, weight, comfort, and cost. While functional near-infrared spectroscopy (fNIRS) instruments have been around since the early 1990s [13], incorporating high-density optode arrays for tomographic reconstruction has dramatically transformed imaging performance [14–16]. Through these advances, high-density diffuse optical tomography (HD-DOT) has become a powerful approach for cortical functional mapping, including naturalistic movie encoding and decoding, with extensive validation against fMRI [4, 7, 15, 17–20]. While these systems support an open scanning environment, they rely on large numbers of optical fibers tethered to bulky server racks. We developed and validated an untethered, whole-head wearable HD-DOT (WHD-DOT) system that matches the imaging performance of fiber-based HD-DOT while enabling high-density optical neuroimaging beyond the laboratory.

Design tensions in wearable DOT systems mirror consumer wearables, where hardware performance, weight, and battery life must be balanced. High-density optode arrays that drive imaging performance increase the weight on the head, complexity of electrical design, and power demands. For example, distributing embedded computing across the cap enables data compression and simplifies control lines, but requires an increase in power consumption. Separately, the field has developed varying approaches for high-density geometries, ranging from flexible, customizable layouts [9, 21] to hexagonal [10] and rectangular arrays [12], each reflecting different trade-offs among spatial sampling, coverage, and mechanical practicality. Finally, all these aspects work against power and weight, which together with battery capacity, constrain imaging duration.

The WHD-DOT system developed here addresses these challenges by combining tomographic imaging with dense spatial sampling, distributed embedded computing, wireless data streaming, and untethered lightweight battery-powered mobility. To evaluate WHD-DOT, we followed a three-fold approach using (1) a set of standard functional localizer tasks, (2) naturalistic movie watching, and (3) skilled piano performance, using repeatability, encoding, and decoding analyses. While the first two approaches follow well-validated paradigms, piano performance shows that complex brain dynamics during free, real-world behavior can be decoded with an unconstrained wearable device.

## Results

### High Performance Wearable Brain Imaging System

Wearable DOT systems are complex, involving >100 optodes, each of which must be positioned in an array and spring-loaded. To manage this complexity, we designed the WHD-DOT system around source-detector (SD) modules, each with two sources and two detectors (**Fig. 1a–b**). These SD-modules require control to encode the sources, and processing to decode the data streaming from the detectors into SD-pair data structures. To accomplish this, digital signal processors (DSPs) are integrated into each SD-module and synchronized across the cap (**Fig. 1c**). To channel light to and from the scalp and provide a combing action through the hair, both the LED sources and photodiode (PD) detectors are coupled to spring-loaded cylindrical light pipes (NA = 0.5, 3 mm diameter, 12 mm length). The SD-modules plug into a wiring harness embedded in a 3D-printed flexible cap substrate, enabling two hours of unconstrained neuroimaging, powered by a battery via a backpack-mounted power supply and wireless data streaming (32.8W power consumption; **Fig. 1d**). The imaging cap contains 64 SD-modules (17g each), housing 128 sources and 128 detectors distributed across the head to provide regularly spaced, high-density cortical sampling (13 mm inter-optode spacing, both within and between modules) and uniform sensitivity coverage (**Fig. 1e–f**).

**Figure 1.**
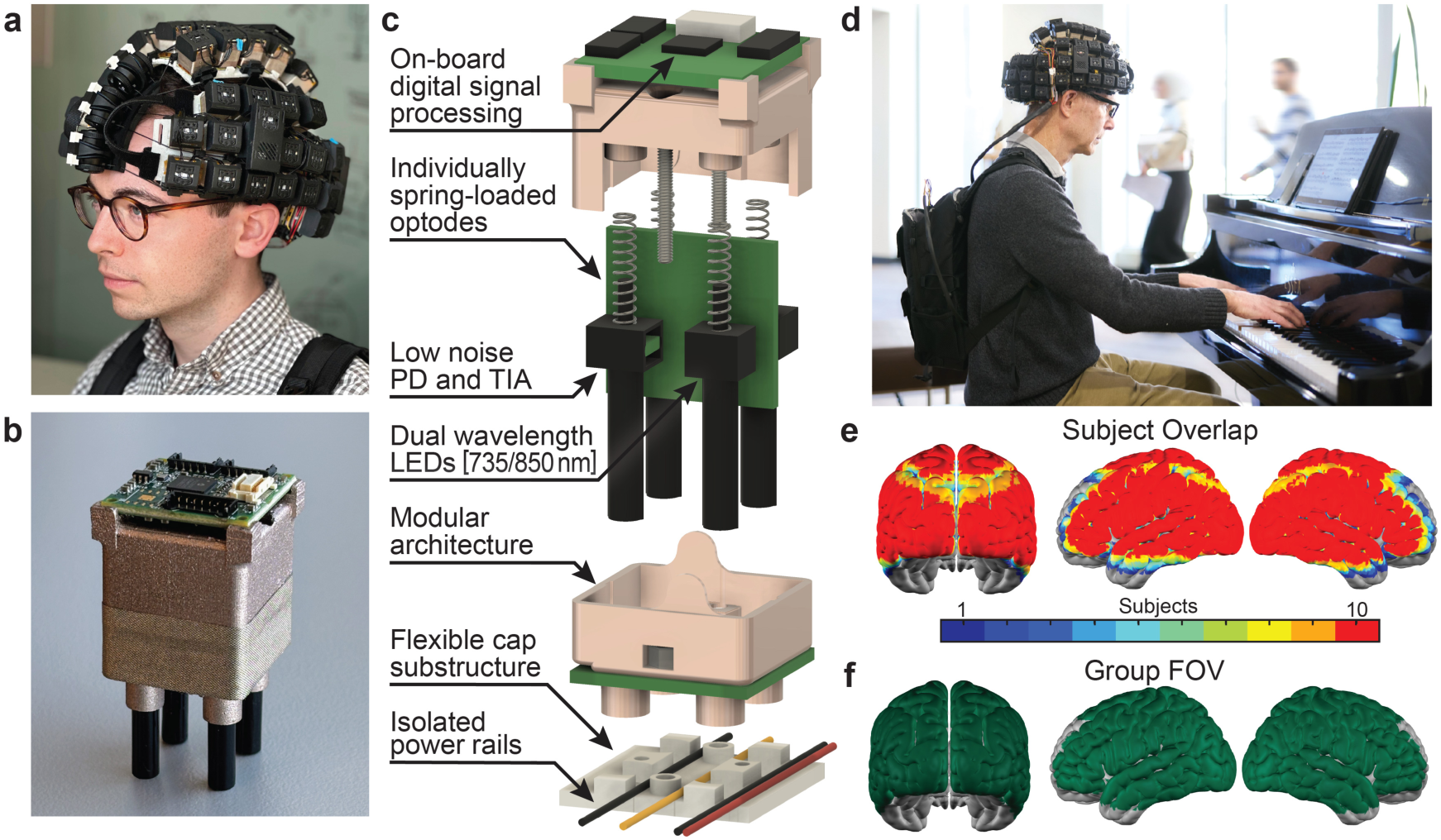
Mechanical and electrical design of WHD-DOT. **(a)** *Pictured: Author W.T.H.* The fully wearable, untethered wearable high-density diffuse optical tomography (WHD- DOT) system enables unconstrained neuroimaging, powered by a battery via a backpack-mounted power supply and wireless data streaming. The imaging cap contains 64 modular source-detector units (SD-modules) distributed across the head to provide regular high-density cortical sampling and uniform sensitivity coverage. The cap provides a whole-head field of view (FOV), including auditory, visual, and motor regions, and is secured with an adjustable forehead ratchet system. Data are transmitted via four integrated Wi-Fi modules embedded within the cap. **(b)** Individual SD-module and **(c)** internal mechanical architecture showing integrated optical and electronic components. **(d)** *Pictured: Author E.J.R.* The untethered system can be used during naturalistic behavior (live piano performance), enabled by backpack-mounted power and wireless data transmission to a nearby acquisition computer. Photo by Matt Miller/WashU Medicine. **(e)** Overlap of subject-specific FOV across participants, demonstrating consistent whole-head cortical coverage (N=10). Sensitivity maps were thresholded at 5% of the maximum sensitivity. **(f)** Group FOV showing the union of sensitivity across participants.

The highest-performing HD-DOT systems have relied on fiber-based architectures, where large cable bundles and rack-mounted electronics deliver the low-noise performance and measurement density required for tomographic imaging [4, 15–17]. Translating this performance to a wearable form factor requires careful noise isolation to prevent high-power digital circuits from interfering with low-noise, high-dynamic-range analog detectors. Optimizations included embedding the transimpedance amplifier (TIA) into the detector circuitry, isolating analog and digital circuits, and shielding modules (**Fig. 1b–c**), leading to a detectivity matching fiber-based systems (**Supplemental Table 1**). This led to weight savings, bringing the whole-head HD field-of-view (FOV) system under 2 kg while maintaining high performance. We designed scalable, distributed, digital control boards using a DSP to simplify cabling, control sources, and compress data for wireless transmission by 96%. The DSP implementation uses a configurable source encoding scheme, supporting spatial, frequency, and temporal multiplexing (**Fig. 2a–b**), to maximize signal-to-noise ratio (SNR) at an 8 Hz whole-FOV frame rate (**Fig. 2c–e**). This allows six dual-wavelength sources to be active simultaneously, with each detector measuring all nearby sources, resulting in a 12x increase in integration time. This source pattern was designed to maximize framerate while minimizing crosstalk between measurements (**Fig. 2f–h**). To capture both short/bright and long/dim channels, a two-pass encoding scheme alternates sources between 50% and 1% duty cycles, providing a >30dB increase in effective dynamic range, to 151dB (**Fig. 2i, Supplemental Table 1**).

**Figure 2.**
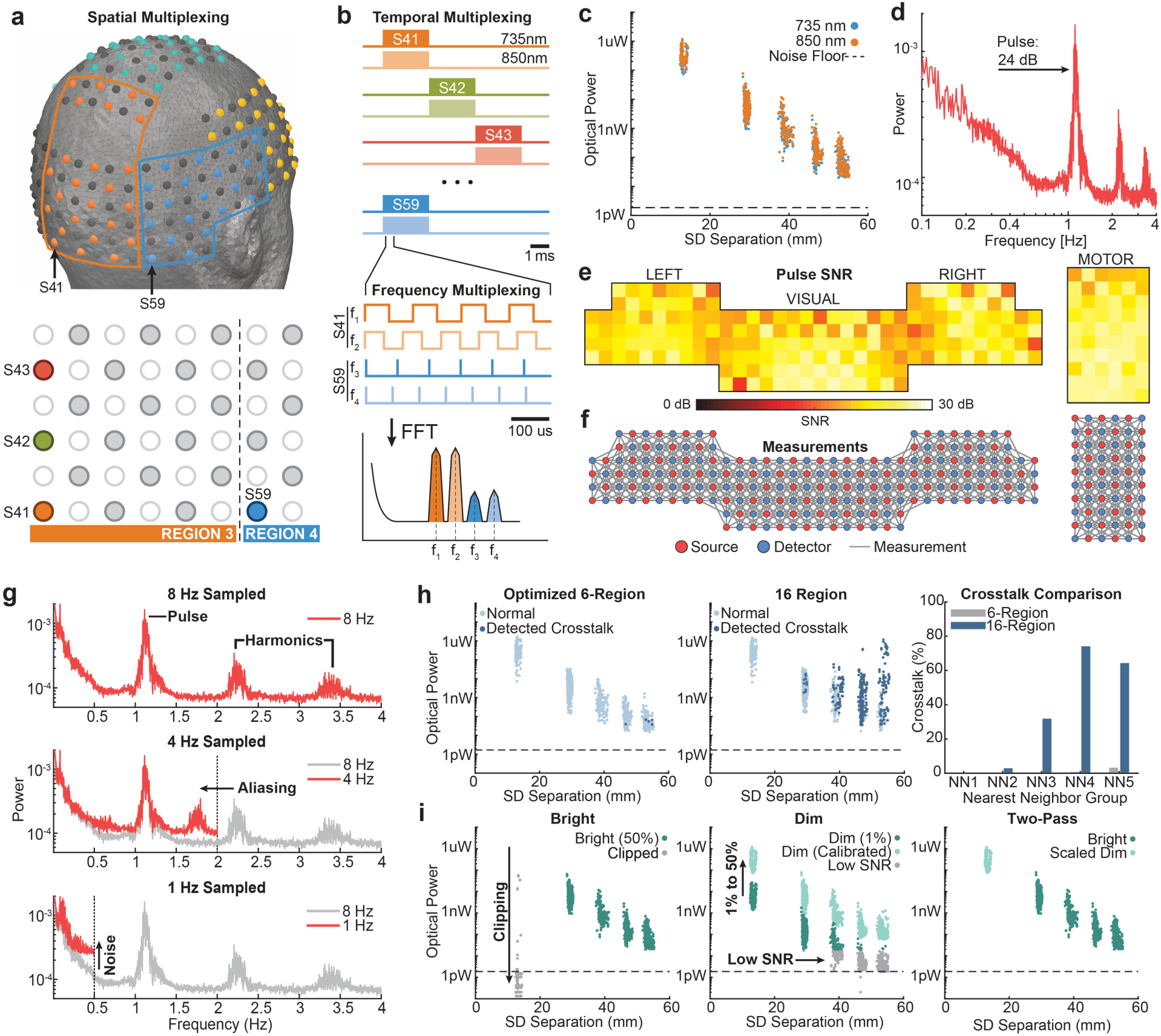
Optical and acquisition performance characteristics of the WHD-DOT system. **(a)** The cap is partitioned into independent source regions, enabling simultaneous illumination while minimizing optical crosstalk and maintaining measurement fidelity. **(b)** The source encoding scheme uses spatial, temporal, and frequency multiplexing to increase sampling efficiency and integration time. Duty-cycle modulation further increases effective dynamic range. **(c)** Dual-wavelength operation (735 and 850 nm) showing the expected log-linear decrease in mean detected light intensity with increasing source-detector (SD) separation, consistent with diffusive light propagation in tissue. **(d)** Frequency-domain spectra demonstrating robust physiological signal capture, including a clear cardiac pulsation peak. **(e)** Aggregated pulse amplitude across optodes can be used as a proxy for cap fit quality and optode-scalp coupling. Pulse signal-to-noise ratio (SNR) is spatially uniform across the cap. **(f)** The regularly spaced source-detector array enables high-density sampling required for tomographic reconstruction, providing 2,976 measurement channels with source-detector separations ≤40 mm. Measurements are densely overlapping, with first nearest-neighbor spacing of 13 mm. **(g)** At 8 Hz, physiological signals are well-separated from functional signals, but lower framerates cause aliasing into the measurement band. **(h)** Optimized 6 region source pattern offers a balance of high framerate and integration time with minimal crosstalk up to 50 mm source-detector separation. Increasing the frame rate by using 16 simultaneous sources dramatically increases crosstalk. (**i**) Two-pass encoding scheme combines 50% and 1% duty cycle passes to extend effective dynamic range by >30 dB. Data for panels **(c–i)** were from a single participant (S11).

An optimal imaging cap must balance lateral rigidity, to keep the SD-modules perpendicular to the local scalp surface, with sufficient flexibility to conform to different head shapes. To accomplish this, we 3D-printed the cap from Thermoplastic Polyurethane (TPU) 70-A, combining stiffness and elasticity. Spring-loaded optodes accommodate further head-shape variations and comb through hair to maintain consistent scalp contact [4]. The optode geometry yielded 2,976 measurements with SD distance ≤40 mm. Participants retained 2,258 ± 77 (mean ± Standard Error of the Mean (SEM)) after temporal variance thresholding (<7.5%; **Fig. S1a**). To evaluate the success of cap fit, we analyzed the pulse SNR, which across retained channels was 10.8 ± 0.9 dB (mean ± SEM, **S1b–c**), confirming consistent optode-scalp coupling.

To perform unconstrained naturalistic imaging outside lab environments, the system must be untethered and tolerate motion. We created a wireless data streaming system with four onboard Wi-Fi modules that transmit data to a nearby laptop, thereby eliminating movement restrictions and motion artifacts caused by fiber bundles and data cables [12]. To track motion across runs, we computed global variance of temporal derivatives (GVTD), a motion detection metric for optical neuroimaging data [22]. Robustness to motion was demonstrated for all participants, with minimal motion artifacts (0.48% ± 0.18% mean ± SEM of timepoints above the GVTD threshold; **Fig. S1d–e**). Live monitoring of data quality metrics, including mean light levels, pulse SNR, and light fall-off curves was enabled through Wi-Fi streaming and a custom real-time MATLAB application.

### Repeatable and localized single-trial functional mapping

To test whether WHD-DOT supports robust single-trial functional mapping, we administered functional localizer tasks, including auditory, visual, motor, and language [15, 17], and performed repeatability and decoding analyses. Single-participant visual block-average maps were localized to visual cortex and spatially similar to subject-matched fMRI maps (**Fig. 3a**). Corresponding single trials for the visual left condition showed matching localized responses (**Fig. 3b**). To test whether the DOT signal contained task-specific activations, a maximum correlation template-matching decoder was applied (**Fig. 3c–d**). Task blocks were divided into training, used to construct task templates, and test, evaluated individually against those templates. To get stable estimates of decoding performance, cross-validation was applied. To assess single-trial performance, test blocks were never averaged. A representative participant achieved well above chance decoding (accuracy = 73.9 ± 3.0%, mean ± SEM, 6-way, chance = 16.7%; **Fig. 3d**). While decoding performance varied across participants, ranging from 32.2% to 73.9% (**Fig. 3e**), all participants performed above chance (accuracy = 52.0 ± 4.3%, mean ± SEM). Data quality metrics (pulse SNR, measurements retained, GVTD) were not indicators for differences in decoding performance (**Figs. S1f–h, S2a**). To assess the robustness of the decoder, we systematically varied the amount of training data used to construct the template. Decoding accuracy continued to improve with additional training data, without reaching a plateau, suggesting that more training data could improve performance (**Fig. 3f**). To isolate the contributions of template and test data, we independently varied one while holding the other constant. Increasing the number of test trials had a marginal effect, whereas increasing the amount of template data led to substantial performance gains (**Fig. S2b**).

**Figure 3.**
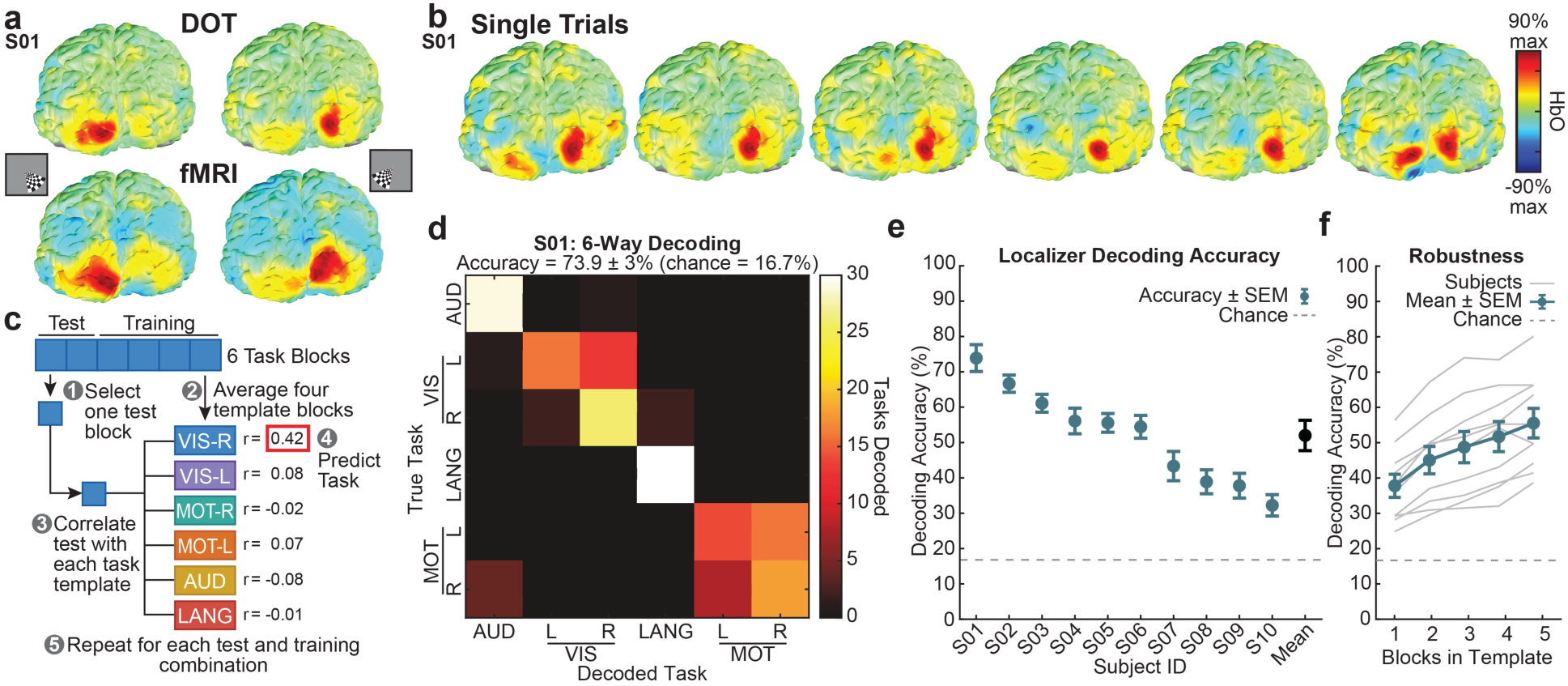
Single-trial repeatability and functional decoding performance (HbO). **(a)** Block average map for a representative visual stimulus run from one participant (S01), demonstrating spatial distribution of trial-to-trial signal repeatability (top). Block average map for the same task and participant in functional magnetic resonance imaging (fMRI; bottom). **(b)** Corresponding single-trial visual left activation responses contributing to the block average maps in **(a)**, illustrating repeatable task-evoked visual responses across trials. **(c)** Schematic of the template-based decoding framework and cross-validation procedure used for localizer classification. A six-way decoder was implemented corresponding to the six task conditions across localizer paradigms: left vs right visual checkerboard stimulation (VIS-L, VIS-R), left vs right finger tapping (MOT-L, MOT-R), auditory word presentation (AUD), and language verb-generation task (LANG). **(d)** Representative confusion matrix from a single participant (S01), showing decoding performance well above chance with partial misclassification between left and right conditions for visual and motor tasks. **(e)** Localizer decoding accuracy across participants sorted by performance. Decoder performance varies across participants, with all participants performing above chance. See **Table 1** for decoding performance across all hemoglobin contrasts: oxy-, deoxy-, total-hemoglobin, and oxy-deoxy difference (HbO, HbR, HbT, and HbDiff, respectively). **(f)** Effect of template averaging on decoding performance. Increasing the number of trials used to construct task templates improves decoding accuracy consistently across participants. Together, these results demonstrate reliable single-trial signals, sufficient to support functional decoding.

**Table 1.** Comparative performance analysis across chromophores and contrasts: oxy-, deoxy-, total-hemoglobin, and oxy-deoxy difference (HbO, HbR, HbT, and HbDiff, respectively; N=10).

| Task | HbO | HbR | HbT | HbDiff |
| --- | --- | --- | --- | --- |
| **Localizer Group Maps** | *max(t)* | *abs(min(t))* | *max(t)* | *max(t)* |
| **VIS (left-right)** | 4.8 | 4.6 | 4.1 | 4.8 |
| **MOT (left-right)** | 1.9 | 2.2 | 2.7 | 2.1 |
| **AUD** | 4.6 | 5.4 | 2.9 | 4.9 |
| **LANG** | 3.3 | 3.3 | 2.9 | 3.4 |
| **Decoding** | *Decoding Accuracy % (Mean ± SEM)* |  |  |  |
| **6-way Localizer** | 52.0 ± 4.3 | 53.3 ± 4.2 | 45.8 ± 3.2 | 54.1 ± 4.5 |
| **6-way Movie** | 48.3 ± 5.2 | 40.0 ± 6.2 | 35.0 ± 6.3 | 46.7 ± 7.4 |

Functional localizer task produced well-localized cortical activation patterns on a group level (**Fig. 4a–d**). To confirm stimulus-evoked hemodynamic responses, oxy-, deoxy-, and total-hemoglobin (HbO, HbR, and HbT, respectively) time courses were extracted from task-specific regions of interest (ROIs) averaged across participants (**Fig. 4e**). All tasks showed the expected hemodynamic responses, with increases in HbO and decreases in HbR following stimulus onset. To validate the anatomical specificity, the WHD-DOT group maps were compared with canonical fMRI maps (**Fig. 4f–h**; N = 15 fMRI reference data, see **Supplemental Table 2**). Activation patterns showed strong spatial correspondence, confirmed by positive correlations between DOT and fMRI maps for matched (mean r = 0.45) and negative-to-low correlations for mismatched tasks (mean r = 0.01; **Fig. 4f**). Subject-level maps also showed strong, localized activations in corresponding sensory regions (**Fig. S2c–f**).

**Figure 4.**
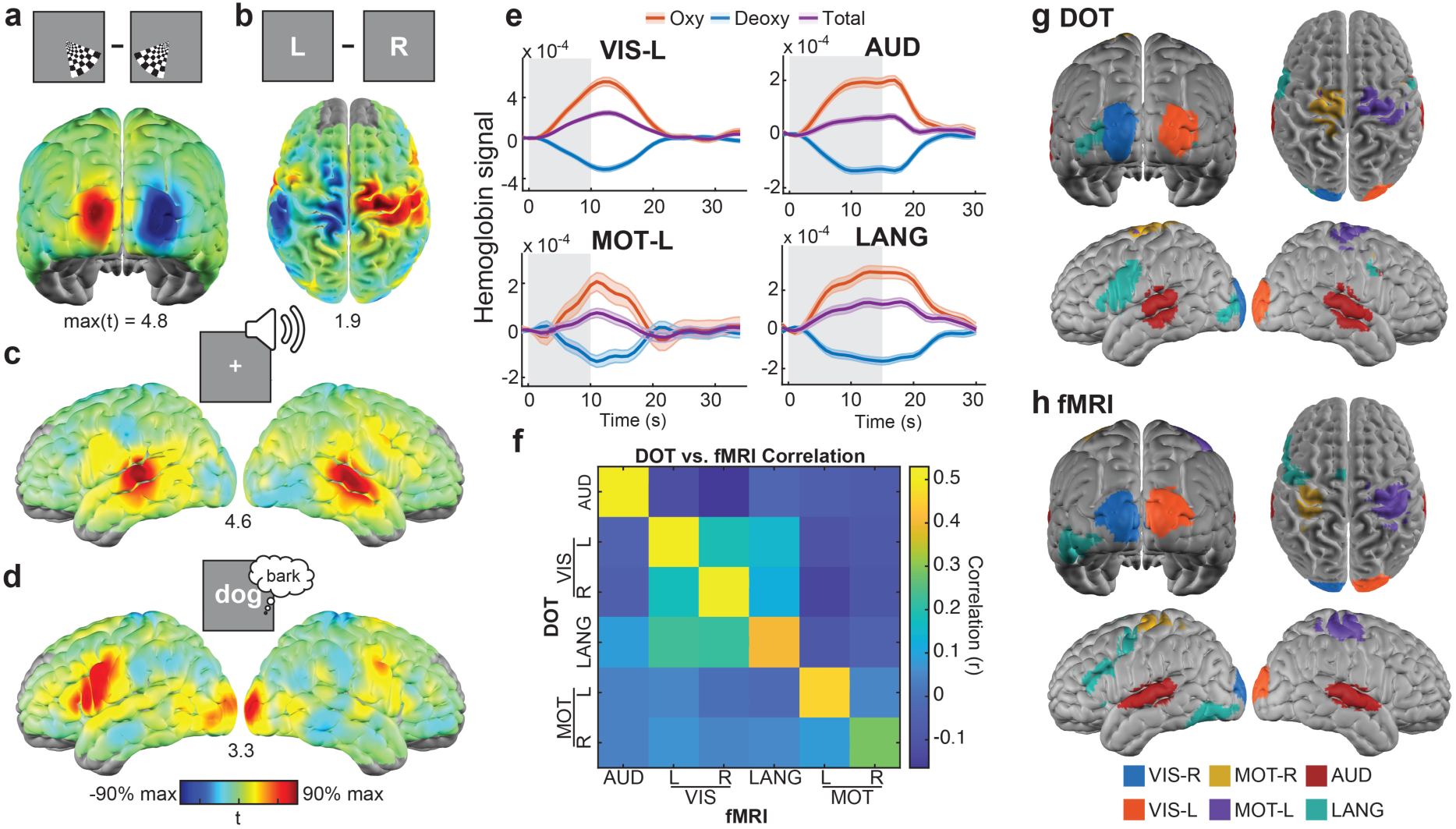
Group-level functional mapping fidelity using wearable high-density optical neuroimaging. (a–d) Group fixed-effect t-statistic activation maps (HbO, N=10) in response to standard functional localizer tasks: **(a)** right vs left visual checkerboard stimulation (VIS-R, VIS-L), **(b)** left vs right finger tapping (MOT-L, MOT-R), **(c)** auditory word presentation (AUD), and **(d)** language verb-generation task (LANG). Numbers below each map indicate the maximum t-value for the corresponding group fixed-effects map. Group maps include all available data from each participant. **(e)** Group-averaged hemodynamic time courses (HbO, HbR, HbT) extracted from task-specific regions of interest (ROIs). HbO shows positive responses, HbR shows negative responses, and HbT shows smaller positive responses consistent with expected neurovascular coupling. ROIs were defined by thresholding each task t-map at 50% of its maximum value. For motor and language tasks, ROI selection was constrained to task-relevant cortical regions to minimize contributions from stimulus-driven visual responses. **(f)** To quantitatively assess similarity between DOT and fMRI localizer group maps, spatial correlation coefficients were computed between matched and mismatched task t-maps (no threshold was applied). **(g–h)** Thresholded WHD-DOT activation maps (HbO) compared to corresponding fMRI group activation maps acquired using the same task paradigms. **(f– h)** fMRI maps were masked by the WHD-DOT FOV, and motor comparisons were restricted to the FOV of the WHD-DOT motor panel for DOT and fMRI. fMRI maps were derived from independently acquired group datasets and are not subject-matched. See **Supplemental Table 2** for details of datasets contributing to group maps.

To evaluate performance across hemoglobin contrasts, we conducted localizer mapping and decoding analyses using HbR, HbT, and oxy-deoxy difference signals (HbDiff; **Table 1**). Group-level HbR activation maps showed spatial localization comparable to HbO- derived maps across tasks (**Fig. S3a–d**). HbR and HbDiff yielded similar accuracy for localizer decoding, with HbT performing lower (**Figs. S3e–g**).

### Distributed multi-modal cortical maps of naturalistic movie viewing

To evaluate whether WHD-DOT can resolve distributed brain responses to naturalistic stimuli, participants viewed a 10-minute audiovisual movie clip (*Despicable Me*) twice. Semantic features were analyzed using manually coded speech and face regressors, which were correlated with the time courses. Speech elicited bilateral auditory and temporal lobe responses (**Fig. 5a**), whereas faces produced strong occipital and ventral visual activations with additional responses in the superior temporal sulcus (**Fig. 5b**). Maps matched spatial patterns from prior HD-DOT movie studies [4, 17]. To demonstrate temporal alignment between features and responses, time courses were extracted from an auditory seed and compared to the speech regressor (r = 0.57; **Fig. 5c**). Similarly, a visual seed was correlated with the face regressor (r = 0.37; **Fig. 5d**), confirming feature- synchronized responses. Results without temporal smoothing yielded comparable spatial patterns and modestly lower seed correlations (Speech = 0.50, Faces = 0.35; **Fig. S4a– d**).

**Figure 5.**
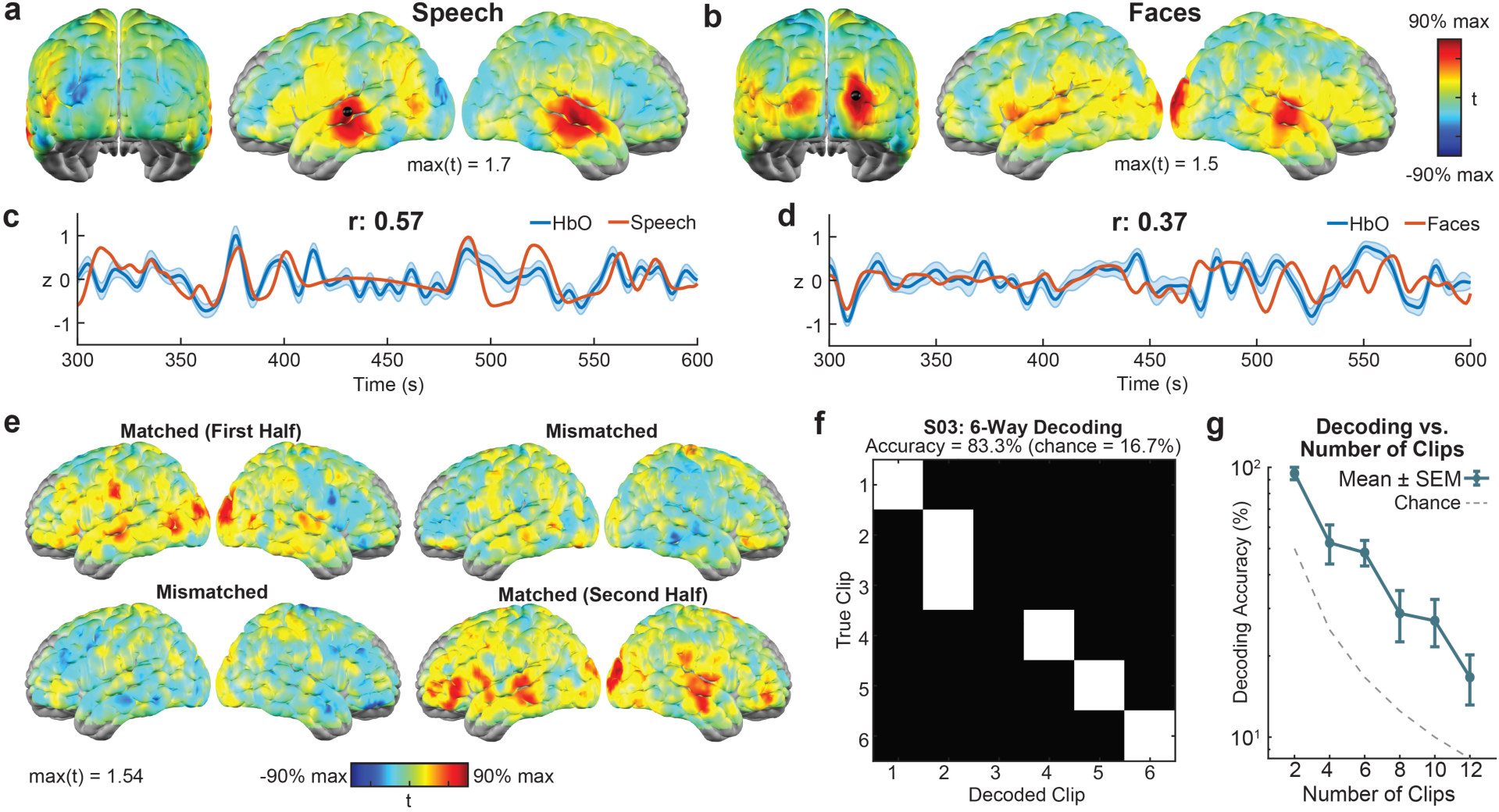
Naturalistic movie feature mapping, cortical synchronization, and stimulus decoding (HbO, N=10). (a–b) Participants viewed a 10-minute audiovisual movie clip twice within a single imaging session. Regressor-based analyses of semantic features, including speech and faces, revealed distinct cortical activation patterns in temporal and visual regions. **(c–d)** Representative voxel time courses showing alignment between cortical activity and stimulus features. Signals from an auditory voxel correlated with the speech regressor (r = 0.57) **(c)**, and signals from a visual voxel correlated with the face regressor (r = 0.37) **(d)**. For visualization, DOT responses were low-pass filtered at 0.1 Hz to match the temporal smoothness of the regressors **(a–d)**. Analyses using unfiltered data are provided in **Supplemental Fig. S4**. **(e)** Voxelwise temporal correlation maps comparing matched and mismatched movie segments across participants (group t-statistic maps). Matched segments (same clip across viewings) show strong cortical synchronization, whereas mismatched segments (cross-clip comparisons) show reduced or negative correlations across the field of view. For this analysis, each movie run was split into two 5-minute clip segments. **(f)** Template-based movie decoding using a maximum-correlation decoder based on spatiotemporal activation patterns across movie segments for a single participant. To separate training and testing data, the first viewing served as the template, and the second viewing as the test dataset. Decoded clips are indicated in white. Decoding performance was well above chance for a six-way decoding task (six 100-second clip segments), indicating that WHD-DOT captures stimulus-specific cortical dynamics. Initial transient periods were excluded from analysis. **(g)** Decoding performance remains above chance across varying movie segment durations on a group level.

To assess the extent of synchronized responses across repeated movie presentations, we performed correlation analyses. Matched clips showed high synchronization across visual, temporal, and frontal regions, whereas mismatched clips exhibited low-to-negative correlations (**Fig. 5e, Fig. S4e–g**). At the single-voxel level, time courses showed similar patterns, with high temporal correlation for matched (r = 0.58) and low correlation for mismatched responses (r = -0.03, **Fig. S4h–i**).

To assess the sensitivity and reproducibility of movie clip-specific cortical dynamics, a spatiotemporal template-matching clip decoder was applied. Decoding performance was high for a representative participant with 83.3% accuracy for a six-way decoding task (chance = 16.7%; **Fig. 5f**). Decoding performance varied across participants, but group-average decoding performance stayed above chance (accuracy = 48.3 ± 5.2%, mean ± SEM, **Table 1**). The robustness of the decoder was tested by varying the number of clips from 2 to 12. Decoding accuracy remained above chance across all clip set sizes (**Fig. 5g**). To compare performance across hemoglobin contrasts, 6-way movie decoding was performed for HbR, HbT, and HbDiff, in addition to HbO. Performance was lower for these contrasts but remained above chance (HbR: 40 ± 6.2%, HbT: 35 ± 6.3%, HbDiff: 46.7 ± 7.4%; mean ± SEM; **Table 1**). These results demonstrate that WHD-DOT captures distributed and repeatable cortical dynamics through encoding and decoding of naturalistic movies.

### Decoding of unconstrained piano play

Piano playing engages a distributed network of auditory, visual, motor, and cognitive control systems and requires continuous bimanual movement, making it a rigorous test of real-world neuroimaging [23–25]. A trained pianist performed continuous piano play across three sessions while wearing the fully untethered WHD-DOT system, enabling natural posture and free arm and hand movement (**Figs. 1d,6a**). Imaging sessions consisted of three 15-minute runs. In each run, the pianist played three ragtime songs consisting of repeating segments, yielding nine repeats of each song across sessions (**Figs. 6b**).

**Figure 6.**
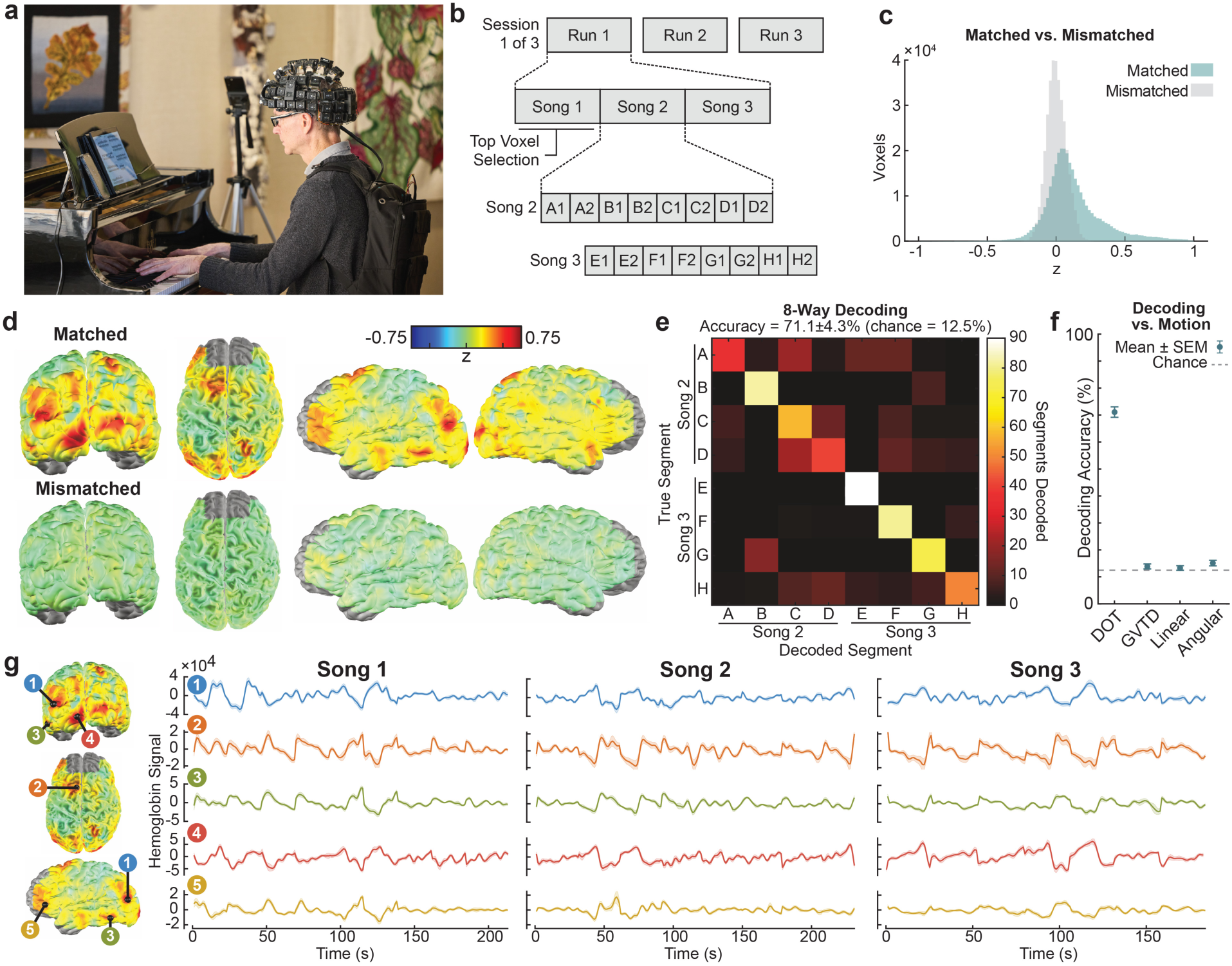
Unconstrained imaging of naturalistic piano playing. **(a)** *Pictured: Author E.J.R.* A skilled pianist played three songs in three imaging sessions while being imaged with WHD-DOT. Photo by Matt Miller/WashU Medicine. **(b)** To test for song-specific information, a decoding task was performed using three ragtime songs, selected for their internal repeating structure. The first song was used to select voxels sensitive to piano playing, and the second two were analyzed for decoding. The songs have four repeating themes, resulting in eight segment types between the two decoded songs. Songs are repeated three times. Within each song, each segment is repeated twice, yielding six repeats per session. **(c)** The histogram of voxel correlations for matched songs shows rightward skew relative to mismatched songs. **(d)** Matched song repeatability maps show distributed positive correlation across the cortex, in brain networks associated with music performance, while mismatched song maps show negative-to-low correlations. (**e**) Cross-validation was performed for all test/template combinations, resulting in a decoding performance of 71.1%, well above chance of 12.5%. **(f)** To test whether motion was a contributing driver of decoding performance, we generated templates from motion metrics. Global variance of temporal derivatives (GVTD, data derived) and linear acceleration/angular rotation from an accelerometer placed on the cap were unable to reliably decode test data. There is a strong separation between the decoding performance of functional DOT and all the motion metrics (which barely exceed chance). **(g)** Time traces of five seeds in multiple cortical locations (averaged across time, shading for SEM) show distinct temporal structure.

To test whether the DOT signal captures song-synchronized brain responses, repeatability was assessed using spatiotemporal correlation analysis across repeated trials (**Figs. 6c–d**). Networks associated with piano playing and music performance, including primary visual cortex, ventral and dorsal visual streams, parietal and frontal areas [23–27], showed stronger correlations, with strongest in visual regions (**Fig. 6d**). Correlation between matching repeated song segments showed a strong skew towards positive correlations (z = 0.127), while correlations of mismatched segments showed a symmetric distribution centered near zero (z = 0.004; **Fig. 6c**). To test whether the DOT signals captured song-specific information, a decoding task was applied. To constrain the analysis to informative voxels, decoding was restricted to the top 10% of voxels selected based on repeatability in a held-out dataset (Song 1). Decoding performance was well above chance, with 71.1 ± 4.3% accuracy (8-way decoding, chance = 12.5%; **Fig. 6e**). To test decoder robustness, the threshold for informative voxels was varied from all voxels (100%) to the top 1% (**Fig. S5a**). Accuracy increased with progressively stricter voxel thresholds, peaking at 5% of retained voxels (74.7 ± 4.9%). Decoding performance benefited primarily from additional template data, consistent with localizer results (**Fig. S5b–d**).

To test for motion confounds, we collected three motion metrics (GVTD, and two derived from a cap-mounted accelerometer) and used these as templates to decode the DOT responses. Decoding performance was close to chance (12%–16%), suggesting decoding is not driven by motion during play (**Fig. 6f**). To assess the temporal dynamics of the piano-related brain responses, five seeds in networks associated with piano performance were identified, and time courses were visualized for the three songs (**Fig. 6g**). The seed time courses showed varying but highly repeatable temporal components across songs, with low correlation to motion metrics (**Fig. S5e)**. Together, these results demonstrate that WHD-DOT can capture distributed, reproducible, and song-specific brain dynamics during unconstrained real-world performance, establishing WHD-DOT for imaging complex multisensory cognitive tasks outside the laboratory.

## Discussion

We developed a wearable, untethered, full-head HD-DOT system and characterized it across four areas: (1) a modular industrial design providing flexibility and stability across many head shapes (**Fig. 1**), (2) an electro-optical design matching fiber-based system performance in a wearable form factor (**Fig. 2**), (3) robust encoding/decoding of functional localizers and naturalistic movies (**Figs. 3–5**), and (4) a real-world naturalistic piano-playing task with reliable repeatability across segments and songs (**Fig. 6**).

A requirement for high-quality optical neuroimaging is maintaining consistent optode-scalp contact across varying head shapes. Hence, cap designs remain an active engineering challenge, with approaches spanning from soft, stretchy caps that conform to individual head shapes [9, 10, 21] to rigid helmets that prioritize stability [11]. We pursued a semi-rigid 3D-printed TPU cap with spring-loaded optodes, balancing stability with the flexibility needed to accommodate multiple head shapes without significant stretching. To achieve a high-density array in a wearable form factor, we designed SD-units with focus on modularity. Each SD-module holds four optodes, maintaining a 13 mm nearest-neighbor spacing both within and between modules. This optimized high-density full-head design supported robust distributed brain mapping in 10 participants using functional localizers and naturalistic movies (**Figs. 3–5, S2–5**).

To achieve high-quality tomographic reconstructions, DOT systems require a high frame rate, dynamic range, and SNR. Achieving this in a wearable form factor requires careful optimization at every stage of the detection pipeline. Detectors must have sufficient sensitivity near their fundamental noise limit to capture long/dim measurements while maintaining the wide dynamic range needed for short/bright measurements. Source encoding requires individual control of timing, frequency, and duty cycle of sources, without increasing the footprint of the system. DSPs need to perform high-speed acquisition of analog measurements and demodulate them at a data rate compatible with wireless transmission. WHD-DOT achieves the same performance characteristics as fiber systems in a wearable form factor, through careful analog design and a distributed, scalable digital control system. Dynamic range, framerate, and even detectivity match those of fiber systems, using the larger optode area to offset a higher noise-equivalent power (NEP; **Supplemental Table 1,** [12]). These system-level benchmarks translate directly into improved imaging performance, evidenced by robust single-trial localizer repeatability (**Figs. 3–4**, **Table 1**).

Wearable neuroimaging systems span a broad range of modalities, each with distinct tradeoffs between spatial resolution, coverage, and wearability. fNIRS offers lightweight, portable solutions but remains limited in cortical coverage and spatial resolution. Emerging wearable, higher-density optical neuroimaging systems have begun to address these limitations [9–12], with several demonstrating tomographic reconstruction [9, 10, 12], though array geometry and imaging FOV vary considerably across systems. With WHD-DOT, we addressed these limitations by providing a full-head FOV through a regularly spaced, high-density array that combines the spatial fidelity of the highest-performing fiber-based HD-DOT systems [4, 15–17] with the mobility of fully untethered wearable systems. The system here improves on an earlier prototype [12] with lighter modules (37% weight reduction), full-head rather than partial FOV (64 instead of 49 modules), and better optimized encoding patterns that enable full-head imaging. To allow for longer continuous imaging sessions, we lowered the power consumption (19% reduction) and doubled the Wi-Fi transmission speed. These improvements were required to enable the fundamental imaging performance and the naturalistic tasks demonstrated.

A key benchmark of any neuroimaging system is the ability to map both localized and distributed cortical activations with sufficient fidelity for single-trial analysis. System performance was rigorously tested with encoding and decoding analyses of functional localizers and naturalistic movies. While encoding analyses provide spatial maps of brain function, decoding assesses whether the functional brain responses contain sufficient information to discriminate stimulus conditions. WHD-DOT showed robust and repeatable functional localizer maps at the group (**Fig. 4**), single-subject (**Fig. S2c–f**), and single-trial level (**Fig. 3a–b**). Above-chance decoding for both localizers (**Figs. 3d–f**) and movies (**Figs. 5f–g**) for multiple hemoglobin contrasts (**Table 1, S3e–g**) confirmed that WHD- DOT detects stimulus-related content at the single-trial level.

Capturing the representations in the brain of real-world multi-sensory cognitive tasks requires an imaging system that imposes minimal constraints on natural behavior. Piano playing is a complex task that integrates multimodal perception, motor control, and higher cognitive planning, requiring years of practice. MRI and positron emission tomography (PET) music encoding studies have required participants to lie still on their backs in a scanner, often relying on imagined performance or simplified scanner-compatible keyboards. MR scanner noise necessitates choosing between audio-free playing or modified sequences for quieter but non-continuous sampling [23–25, 28]. Alternatively, sparse fNIRS (<50 measurements [29]) has enabled studies of free piano playing, but largely relies on listening or simplified block designs rather than active, continuous playing or music decoding. By overcoming both the physical constraints of MR scanners and the sparse measurements of fNIRS, our fully wearable high-density system with a full-head FOV established decoding of continuous live piano play (**Fig. 6, S5**). Using repeated song segments, we showed high repeatability in brain regions that correspond to cortical networks implicated in music perception and performance, including visual, parietal, and frontal areas [23–27]. Both time courses and decoding performance were robust to motion confounds, confirmed by data-derived and data-independent motion metrics (**Fig. 6g, S5e**).

Decoding performance varied across participants for both functional localizers and naturalistic movies but was not significantly related to differences in data quality measures, including GVTD, SNR, and measurements retained (**Fig. S1f–h**). Intersubject variability is consistent with prior DOT [7, 17, 20] and fMRI [2, 30] studies. Increasing the number of trials used to construct templates improved accuracy without reaching a performance plateau (**Figs. 3f, S2b**), suggesting that additional data would improve decoding accuracy. Consistent with this hypothesis, prior HD-DOT studies using datasets with more repeats or longer recordings have reported higher decoding performance [7, 18, 20, 31].

WHD-DOT opens several future directions, including multimodal integration, clinical translation, and real-world decoding. First, concurrent measures of behavior and physiology, such as eye tracking, respiration, and cardiac signals, would enrich the interpretation of brain responses during naturalistic tasks. Second, clinical and pediatric populations, including individuals with implants [32], children with ASD [33], or patients requiring bedside monitoring [34] would particularly benefit from wearable optical neuroimaging. While currently optimized for adults, reducing the cap size and weight would enable pediatric imaging with WHD-DOT. Finally, extending continuous language decoding to real-world environments would unlock practical applications for augmented communication [3, 35]. Recent work has demonstrated the feasibility of advanced semantic mapping approaches with a fiber-based HD-DOT system [20], setting the stage for semantic mapping in a wearable system, outside the lab environment.

This wearable, untethered, full-head WHD-DOT system represents a critical step toward high-fidelity neuroimaging of complex behavior and brain function in real-world environments.

## Methods

### Experimental Model and Study Participant Details

#### Participants

Data were collected from 11 participants (32 ± 11 years old, mean ± standard deviation, 6 female). Ten participants were recruited for the functional localizer and movie paradigms, and one participant was recruited for the piano play. All were healthy and had normal or corrected-to-normal vision. Informed consent was obtained from all participants. Consent procedures were conducted in accordance with the IRB protocol approved by the Human Research Protection Office at Washington University School of Medicine. Participants were compensated at a rate of $25 per hour. No participants were excluded.

### Method Details

#### WHD-DOT Instrumentation

The source optodes (**Fig. 1c**) house a small PCB with a 735 and 850 nm dual-wavelength LED source (Marubeni SMT735D/850D), each color driven separately by an n-channel MOSFET (Texas Instruments CSG17484F4). The LED sources output 8 mW or 1.14 mW/mm^2^ (lower depending on the encoding pattern), under the ANSI safe limit of 4 mW/mm^2^. Detector optodes use a PIN photodiode (Hamamatsu S16392-01CT), a low-noise transimpedance amplifier using an op-amp (Texas Instruments OPA1652AIDRGT), and a 30 MΩ feedback resistor on the reverse side of the board, providing high gain while minimizing noise from long signal paths. This maximizes gain while allowing enough bandwidth for frequency division multiplexing.

A digital signal processor (DSP) handles source encoding and analog acquisition for two sources and two detectors in each SD-module (**Fig. 1b,c**). Sources in the cap are grouped in six regions, with one source active per region at a time (**Fig. 2a**). Four source modulation frequencies are used (7.9, 10, 12, 14.2 kHz), with each source using one of two frequency pairs (7.9/10 or 12/14.2 kHz), to simultaneously drive its two wavelengths. Adjacent regions use alternating pairs, allowing nearby detectors to make four simultaneous measurements (**Fig. 2b**). To increase dynamic range, this is used in combination with a two-pass encoding scheme, where regions alternate between modulating at 50% (bright, high power) and 1% (dim, low power) duty cycle. Dim pass values were scaled to match the bright pass with the ratio sin(*πD_bri_*_gℎ*t*_) / sin(*πD_dim_*) as the Fourier coefficient for a first harmonic of a square wave with duty cycle D scales with sin(*πD*). Detector signal levels are collected at 250 ksps with a 16-bit ADC through a 4- channel digital isolator to prevent digital noise coupling into the analog detector circuit. The DSP demodulates the acquired signal for each time-encoding step to find the measured power for each of the four frequencies, as well as maximum/minimum voltage to detect clipping.

The cap substructure (**Fig. 1c**) was 3D printed by Protolabs in TPU 70-A, which provides a balance of durability and flexibility. The remaining cap parts, including SD-cube housings tops/bottoms/caps, optodes, supports, cable guides and the ratchet holder were 3D printed in house with FormLabs Black V4 resin.

#### WHD-DOT Encoding Pattern Optimization

Evaluation of frame rate, encoding region crosstalk, and two-pass encoding were derived from several rest runs on a participant. For framerate, an 8 Hz acquisition run was downsampled in a postprocessing step to both 4 and 1 Hz for comparison (**Fig. 2g**). The measurement spectrum was obtained from the FFT of the log-mean normalized time series, averaging across retained second nearest neighbors (∼29 mm) at the 850 nm wavelength.

For encoding regions, four patterns (4, 6, 8, and 16 regions) were evaluated in sequence without moving the cap. The 4-region pattern (which has the smallest number of simultaneous active sources) and the 8-region pattern were used as a baseline for determining crosstalk by taking the minimum light level for measurements in either pattern. Measurements in the 6 or 16-region patterns with a >50% increase in optical power were considered affected by crosstalk (**Fig. 2h**). Analysis was constrained to measurements retained across all runs. Sources in the 6-region pattern were timed and reordered to maximize distance between simultaneous active sources.

To evaluate two-pass encoding, dim and bright passes were individually extracted from the data in a run (**Fig. 2i**). Low SNR measurements were defined as 20 dB over the system noise floor. Clipped measurements in the bright pass were replaced with measurements from the scaled dim pass, scaled by the corresponding duty cycle.

Analysis was constrained to the visual pad in light fall-off plots, and extended to 60 mm to show the decay to the system noise floor (**Fig. 2c,h,i**). Pulse analysis was constrained to the 850 nm wavelength and second nearest-neighbor (∼29 mm) measurements (**Fig. 2d,e,g**).

#### WHD-DOT Experimental Design

Of the eleven participants, ten completed two imaging sessions on separate days. Session 1 comprised six functional localizer tasks, and session 2 comprised naturalistic movie viewing and a subset of functional localizers (visual and auditory). Participants performed the tasks while comfortably sitting on a chair. One participant, a pianist, completed three dedicated piano play sessions on separate days. All participants underwent an MRI session to obtain anatomical imaging for subject-specific light modeling. Here, functional localizer tasks were collected to inform head and light modeling (See section WHD-DOT Image Reconstruction and Spectroscopy). For four participants, only anatomical images were collected. For these individuals, group-average canonical fMRI maps (N=15) were used to inform head and light modeling (**Supplemental Table 2**).

Cap fitting followed established protocols [15, 17], with adjustments specific to this study. For medium to long-haired participants, hair was parted at midline, and four pigtails were placed between the motor and side panels. For long-haired participants, the pigtails were braided. Optodes were combed through hair. Real-time readouts of light level, signal-to-noise ratio, and the number of measurements retained guided the cap-fitting procedure.

Cap fits were kept consistent across imaging sessions using a precision tape alignment method [36]. Before placing the cap, hypoallergenic silicone tape was applied on both sides along a line extending from the tragus to the lateral eyebrow. The cap borders were traced on the tape, which was then preserved. For future sessions, a new tape was made using the saved tapes as a stencil. The new tape was aligned with the facial fiducials, and the cap was realigned with the tape traces and positioned relative to the inion.

To ensure accurate temporal alignment between WHD-DOT recordings and various stimuli, the localizer synch signals, movie audio, and piano audio were recorded directly during imaging via a NI USB-6212 DAQ device at 48 kHz, connected to a computer handling both data and stimulus acquisition. Wireless data and stimulus synchronization was performed by aligning the arrival timestamp of the first Wi-Fi packet with the timestamp of the first stimulus.

#### WHD-DOT Stimuli

Six functional localizer tasks were included in session 1: (1) auditory, (2) language, (3) visual left, (4) visual right, (5) motor left, (6) motor right. In session 2, participants completed the visual and auditory functional localizer tasks and viewed an audiovisual naturalistic movie clip twice to enable repeatability analysis. The functional localizer tasks followed our standard DOT protocols [15, 17]. Unless otherwise noted, participants maintained central fixation on a crosshair throughout all tasks. See **Supplemental Table 2** for the amount of data collected in each participant.

Auditory: Participants passively listened to spoken word lists (1 word/s) during six 15-s stimulus blocks, each separated by 15-s silent rest periods.

Language: Nouns were presented visually (1 word/s), and participants silently generated a corresponding verb for each word. The task consisted of six 15-s stimulus blocks, each followed by 15-s rest.

Visual (left and right): Black and white checkerboard wedges flickered at 8 Hz for 10 s in the lower left or right visual field, followed by 24-s rest, over 12 pseudorandomized blocks (6 per side).

Motor (left and right): Left or right finger tapping was cued by a centrally presented ’L’ or ’R’ for 10-s, followed by 25-s rest, over 12 pseudorandomized blocks (6 per hand). Participants were instructed to remain as still as possible between blocks.

Movie Viewing: Participants watched a 10-minute audiovisual clip from *Despicable Me* without central fixation, presented twice within session 2. This clip has been used in the Healthy Brain Network (HBN) dataset [37] and validated across encoding paradigms [38], providing a well-characterized naturalistic stimulus for comparison across studies.

#### WHD-DOT Piano Play Paradigm

The piano data were collected over three imaging sessions on separate days, with three runs per session, each repeating three songs for a total of nine repeats (**Fig. 6b**). The cap fit was kept consistent through a precision approach (see WHD-DOT Experimental Design). The songs were *Maple Leaf Rag* (Song 1), *Swipesy* (Song 2), and *Pineapple Rag* (Song 3), all by Scott Joplin, played at a quarter note tempo of 70, 55, and 60 bpm, and lasting 248, 319, and 292 seconds, respectively. Each song contained four repeating segments (A, B, C, D) in the repeating pattern AABBACCDD. The third repeating A segment was ignored to retain equal amounts of each segment type. Repeating segment durations were 27.4 s for *Maple Leaf Rag*, 34.9 s for *Swipesy*, and 32.0 s for *Pineapple Rag*. Segment time points were manually coded from an audio-video recording of the imaging session. Only data from the repeating segments were used for decoding and repeatability analysis.

Motion was recorded with a YostLabs 3-Space Data Logger mounted centrally on the motor pad of the cap. Two motion metrics were used from the accelerometer: linear acceleration and angular rotation. Angular rotation was computed as the norm of the temporal derivatives of head orientation [22], and by the same approach, linear acceleration was computed as the norm of the temporal derivatives of the sensor’s gravity corrected acceleration vector. Alignment of the accelerometer data was performed using a video recording of the imaging run and confirmed by cross-correlating GVTD with the two motion metrics. Accelerometer data were acquired at 200 Hz and resampled to 1 Hz to match GVTD and DOT data.

#### MRI data collection and processing

MRI data were collected on a separate day for subject-specific light-modeling for DOT. Data were acquired on a 3T PRISMA Fit scanner using 20- or 64-channel head coils. Each participant underwent T1-weighted MPRAGE (echo time (TE) = 3.13 ms, repetition time (TR) = 2,400 ms, flip angle = 8°, 1 × 1 × 1 mm isotropic voxels) and T2- weighted (TE = 84 ms, flip angle = 120°, 1 × 1 × 1 mm voxels) structural scans, followed by functional localizers (gradient spin-echo EPI; TE = 33 ms, TR = 1,230 ms, flip angle = 63°, 2.4 × 2.4 × 2.4 mm isotropic voxels, multi-band factor = 4).

Data were preprocessed using fMRIPrep 22.0.2 [39], which is based on Nipype 1.8.5 [40]. To match the HD-DOT localizer preprocessing, data were detrended and bandpass filtered (0.02–0.2 Hz) before smoothing with an isotropic Gaussian smoothing kernel (13 mm FWHM). Data were converted to a percent BOLD change measurement by subtracting and dividing by the average BOLD signal in each voxel over time [16]. For comparison with DOT, fMRI data were resampled to 1 Hz. Full preprocessing details are provided in Supplemental Methods.

#### WHD-DOT data pre-processing

Raw detector measurements are demodulated for each four encoding frequencies with the on-board DSP before being transmitted wirelessly to the real-time monitoring application. The application organizes measurements by source-detector pair, and scales them to input referred optical power, including the two-pass duty-cycle adjustment. Clipped bright pass (50% duty cycle) measurements are replaced with scaled dim (1% duty cycle) measurements.

DOT data were processed similarly to previously reported studies [4, 15, 17]. Raw light levels were converted to differential log-mean intensity values. Channels were rejected if their temporal standard deviation exceeded 7.5% of the mean light level [15], indicating contamination by non-physiological variance, such as head motion. Further channel rejection was performed following the methods in [12]. This included unnaturally large swings in magnitude (baseline drift, a log-mean range of more than 0.2 after a 60-s moving average), dead channels (standard deviation less than 10^-9^), or sudden spikes (a log-mean range exceeding 0.75, or a peak-to-peak intensity variation exceeding 60% of the mean). Rejected channels were excluded from image reconstruction for the entire run. Across participants, 1,873 to 2,681 of the 2,976 possible measurement channels with source-detector separations ≤ 40 mm were retained (2,258 ± 77, mean ± SEM; **Fig. S1a**). Cardiac pulse signal-to-noise ratio (SNR) was computed for every run as the ratio of pulse signal power to background noise floor, expressed in decibels (dB) as 10*dB* log_10_ 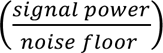. To compute pulse SNR, first an FFT is taken for each retained log- mean measurement, constrained to second nearest-neighbors and the 850 nm wavelength. We then find the maximum frequency bin of each channel’s FFT between 0.85–2.5 Hz, and select the peak pulse frequency as the median across all these values. Signal power is computed as the sum of squared FFT magnitudes within a 1 Hz band centered on the peak pulse frequency. Noise power is estimated as the median squared FFT magnitude of surrounding frequencies (excluding the signal bin and the second harmonic), scaled to the same bandwidth as the signal power. Mean SNR across sessions varied from 7.1 dB to 16.1 dB across participants (10.8 dB ± 0.9 dB, mean ± SEM; **Fig. S1b–c**). To assess head motion per run, the global variance of temporal derivatives (GVTD) was computed for each time point as the root mean square of the temporal derivatives of all measurements [22]. Motion was low across participants and tasks, with a mean of 0.48% ± 0.18% (mean ± SEM) of timepoints exceeding the GVTD threshold of 1.5 × 10^-3^ (**Fig. S1d–e**).

Data were detrended and then lowpass filtered with a 1 Hz cut-off. Superficial signal regression was performed using the average of the first nearest neighbor channels (13 mm separation) to reduce scalp and systemic contributions [41, 42]. Localizer tasks were filtered with low- and high-pass filters at 0.2 Hz and 0.02 Hz, respectively. Movie viewing data were filtered at 0.2 Hz and 0.009 Hz, respectively. Finally, data were resampled to 1 Hz for further analysis.

#### WHD-DOT Image Reconstruction and Spectroscopy

Following best practices for high-performing fiber-based HD-DOT systems, subject-specific light models were generated for each participant from their anatomical MRI scans [4, 17]. Head models were derived from the first imaging session and assumed valid for the subsequent sessions under the precision tape alignment procedure. Optode positions were initially registered to an MRI-derived five-layer segmented head mesh, which accounts for the distinct optical properties of scalp, skull, cerebrospinal fluid (CSF), gray matter (GM), and white matter (WM) [15, 41]. Anatomical landmarks (tragus, inion) were aligned to photographs collected during the imaging session. Segmentation was performed using FreeSurfer [43] and NeuroDOT [15], and the mesh was generated with NIRVIEW [44]. Optode positions were then relaxed onto the head using an iterative optimization that balances source-detector distances with placement on the head mesh [14]. Photon diffusion through the mesh was modeled with NIRFAST’s finite-element solver [45]. Head models were refined by comparing functional DOT and fMRI maps for visual localizer tasks using overlap maps and dice coefficients [17]. Array placement was refit if necessary. This procedure ensured accurate alignment of optodes to subject-specific anatomy and function. DOT images were reconstructed in 2 mm isotropic voxel space, with the Tikhonov regularization constant *λ*_1_ = 0.01, and the spatially invariant regularization parameter *λ*_2_ = 0.1, using subject-specific light models to solve the DOT inverse problem [14, 15, 46]. Relative changes in oxygenated and deoxygenated hemoglobin (HbO, HbR) were derived using spectral decomposition of the absorption changes at 735 and 850 nm [7, 31].

To constrain the analyses to the optical field-of-view (FOV), data were masked at 5% of the maximum of the normalized flat-field reconstruction. All analyses were performed on HbO unless otherwise specified (**Fig. 1e**). The group FOV was defined as the union of sensitivity coverage across the subject-specific FOVs (**Fig. 1f**). Decoding analyses were additionally constrained to a brain mask (CSF, GM, WM).

#### Functional Localizer Analysis

For single-subject and group maps, fMRI and DOT localizer data were analyzed with a general linear model (GLM) in voxel space. Task regressors were convolved with a canonical hemodynamic response function previously derived for HD-DOT [47]. Beta values were estimated for each run, representing stimulus-evoked responses relative to baseline rest periods. Group-level functional localizer maps were derived from fixed-effects t-statistics. For fMRI data, group maps were derived from the canonical fMRI data (N=15; **Supplemental Table 2**).

HbO, HbR, and HbT time courses were extracted from task-specific regions of interest (ROIs) and averaged across participants. ROIs were defined by thresholding DOT t-maps at 50% of the maximum. For time-course analyses, DOT motor and language task data were restricted to the motor and side panels, respectively. Baseline subtraction was performed using the 1s pre-stimulus frame. For comparison with canonical fMRI, the thresholded group t-maps were masked to the WHD-DOT FOV and binarized (1 = inside ROI, 0 = outside ROI). For motor tasks, maps were additionally restricted to the FOV of the WHD-DOT motor panel. To quantify the similarity of task maps, correlation coefficients were computed between DOT and fMRI for all tasks. The correlation analysis was applied to the unthresholded data (not binarized), constrained to the WHD-DOT FOV and motor panel for the motor tasks, and the WHD-DOT FOV for the remaining tasks.

For visualization single subject and group-level maps were projected onto the MNI152 atlas [48, 49] cortical surface, while individual single trial maps were displayed on subject-specific cortical surfaces derived from each participant’s MRI.

Six-way functional localizer decoding was performed using a cross-validated maximum correlation template-matching approach ([17, 31]). Individual task blocks were extracted and averaged across timepoints 5 to 20 seconds post stimulus onset. For visualizing the single trials used in the decoder, this 5 to 20 second block after stimulus onset was averaged. This was compared against fMRI by averaging single trials in the same subject. For decoder cross-validation, in each fold two blocks per task were held out as test data, and the remaining four blocks (training data) were averaged to construct a task template. To test for single trials, decoding was performed on individual blocks (test data were never averaged). Test blocks were correlated with each template, and the template corresponding to the maximum correlation was selected as the predicted label. All unique combinations of four blocks for template construction were exhausted, yielding 15 cross-validation folds (*C*(6,2) = 15). Each fold produced two decoding instances (one per test block), resulting in 30 instances total. Mean decoding accuracy and SEM were computed for each participant and at the group level. Confusion matrices were aggregated at both the single-subject and group level. To assess decoder robustness, the number of blocks assigned to template and test sets was varied jointly and independently. For joint analysis all combinations of template/test blocks were evaluated (five configurations of test/template blocks: 1/5, 2/4, 3/3, 4/2, and 5/1). For independent analysis, six random permutations of test/template blocks were evaluated (varying either test/template from 1 to 5 blocks, while holding template/test constant with 1 block).

Localizer encoding and decoding analyses were compared across hemoglobin contrasts (HbO, HbR, HbT, HbDiff; **Table 1**).

### Naturalistic Movie Encoding Analysis

Movie audio was recorded directly during imaging via the NI DAQ device. Movie onset times were determined by cross-correlating the recorded audio trace with the original movie audio file, providing frame-accurate alignment between brain data and stimulus features.

Two salient naturalistic features were selected for encoding analysis: speech and faces. Binary presence regressors (1 = present, 0 = absent) were manually coded by two independent raters across the full movie duration and averaged to reduce labeling error. Regressors were convolved with a canonical hemodynamic response function, bandpass filtered to match the HD-DOT data (high-pass: 0.009 Hz, low-pass: 0.2 Hz), and z-scored.

For each participant and movie run, voxelwise HD-DOT time series were z-scored across time. Pearson correlations were computed between each regressor and each voxel’s time course, Fisher Z-transformed, and averaged across the two movie runs to yield a single encoding map per participant. Group-level maps were derived using fixed-effects t- statistics. Oxyhemoglobin time courses were extracted from feature-relevant seeds averaged across participants: auditory cortex for speech (MNI: [48, 29, 22]) and visual cortex for faces (MNI: [17, 2, 26]). For regressor-response correlation analyses, voxel time courses were additionally lowpass filtered at 0.1 Hz to match the temporal smoothness of the feature regressors (unfiltered results: **Fig. S4a–d**).

To assess the reproducibility of cortical responses across repeated movie viewings, two complementary repeatability analyses were performed. In the first, the movie was divided into two 300-s segments, and voxelwise temporal correlations were computed between matched clips (same segment across viewings) and mismatched clips (different segments across viewings), yielding four group-level synchronization maps. In the second, full-run correlations were computed between matched viewings and a scrambled version of the second viewing, generated by splitting the run in half and reversing the order of the two halves. Both analyses used z-scored voxel time courses prior to correlation, Fisher Z- transformed correlation coefficients, and fixed-effects t-statistics for group-level maps. Representative oxyhemoglobin time courses were extracted from a visual seed (MNI coordinate: [13, 4, 26]) and averaged across participants (mean ± SEM), with inter-run Pearson correlations quantifying reproducibility.

#### Naturalistic Movie Decoding Analyses

Movie clip decoding was performed using a template-matching approach previously established for HD-DOT [7, 17, 18]. The first movie run served as training data (template) and the second as the test set. The 600-s movie was divided into 2, 4, 6, 8, 10, or 12 segments, corresponding to segment durations of 300, 150, 100, 75, 60, and 50 s, respectively. To minimize HRF carryover from adjacent segments, the first 3 s of each segment were removed. This parameter was optimized in one participant by systematically varying the removal duration from 0 to 10 s and selecting the value that maximized decoding performance. Within each segment, voxel time courses were z-scored across time. Pearson correlations were computed between template and test segments, and the segment with the maximum correlation was selected as the decoded clip. Decoding accuracy was defined as the percentage of correctly identified segments across all test segments. Chance performance was computed as 1/N. Mean ± SEM decoding accuracy across participants was plotted as a function of segment number and compared to chance.

Movie decoding analyses were compared across hemoglobin contrasts (HbO, HbR, HbT, HbDiff; **Table 1**).

#### Piano Data Analysis

To synchronize the multiple instances of piano playing, repeating song segment timepoints were manually coded from an audio-video recording of the imaging session. All piano analysis used the HbO chromophore.

For repeatability analysis, the runs were segmented according to the manually coded time points. Matching block types were trimmed to the minimum block length of that segment type across all runs, then concatenated in time to produce a time-synchronized set of runs. To perform the mismatched repeatability analysis, blocks were randomly ordered before concatenation. Voxelwise temporal correlations were computed between all within-session run pairs, and Fisher r-to-z transformed for visualization. Five seed locations were selected in high-correlation regions across the cortex, and time traces were averaged from concatenated runs.

For piano template decoding, Song 1 was used exclusively to select repeatable voxels. Repeatability was computed the same as the full run maps (concatenated blocks, voxelwise temporal correlation across within-session run pairs), then thresholded to the 10% of most repeatable voxels. Songs 2 and 3 provided eight segment types (four per song) with 6 repeats of each segment type (**Fig. 6b**). For decoding, four blocks were averaged to create a template, leaving two as test. Pearson correlation was performed between template and test, selecting the maximum correlation as the decoded segment type. For cross validation, all 15 combinations of four template blocks were evaluated for each two remaining test blocks, and aggregated across three sessions for 90 total comparisons. Analysis was constrained to subject-specific FOV and masked to the brain volume.

Motion metrics (GVTD, linear acceleration, and angular rotation) were correlated with squared DOT data as the motion metrics are positive semidefinite. The same template/test decoding procedure was then performed between the motion metrics and DOT data. Seeds were similarly correlated with squared DOT data, while using between-run data correlation for DOT.

To assess decoder robustness, we varied the voxel thresholding (1, 2, 5, 10, 20, 50, and 100% of voxels retained, ranked by repeatability computed from Song 1) and the number of blocks in template and tests sets were varied, as with localizer decoding. For joint analysis all combinations of template/test blocks were evaluated. For independent analysis, six random permutations of test/template blocks were evaluated. Decoding accuracy is reported as mean ± SEM across cross-validation folds.

### Statistics and Reproducibility

Unless otherwise indicated, *N* refers to the number of participants (N=11), and all analyses were performed at the individual-subject level before computing group statistics. Sample size was determined based on prior HD-DOT studies employing similar encoding and decoding analyses [12, 15–17], which have reported robust effects at comparable sample sizes.

Encoding and decoding analyses for functional localizers, naturalistic movies, and piano play were quantified using mean and SEM across participants. Group-level maps were derived using fixed-effects t-statistics. Cross-validation was applied to functional localizer and piano decoding analyses to obtain unbiased performance estimates. Movie decoding was performed without cross-validation due to the limited number of movie repeats. Decoding chance level was defined as 1/N, where N is the number of stimulus conditions or movie segments, and served as the reference for evaluating decoding performance. Matched and mismatched repeatability analyses were computed for movies and piano play to establish a null distribution for assessing the content-specificity of cortical responses. For naturalistic movie encoding, speech and face regressors were coded by two independent raters. Inter-rater agreement was quantified using Pearson correlation prior to averaging across raters.

All analyses were performed in MATLAB (version 2020b).

## Resource Availability

### Lead Contact

Requests for further information and resources should be directed to and will be fulfilled by the lead contact, Joseph P. Culver & William T. Hamic.

## Data and code availability

All data will be deposited and are publicly available as of the date of publication. Code for reproducing the results from this paper will be deposited and is publicly available as of the date of publication. Any additional information required to reanalyze the data reported in this paper is available from the lead contact upon request.

## Acknowledgements

This work was funded by the National Institutes of Health [grant numbers U01EB027005, R01NS090874, R01EB034919] awarded to J. P. C., the Washington University’s Imaging Science Pathway Fellowship awarded to W.T.H., W.F., and M.F. [grant number T32EB014855], the CCSN pathway traineeship to H.E.D. [grant number T32NS115672], the SPIE-Franz Hillenkamp Postdoctoral Fellowship awarded to M.F., and the American Heart Association [grant number 26POST1562738] awarded to M.F.. The authors thank all participants for their generous time dedicated to this research.

## Author contributions

System design and building: W.T.H., A.S.A., S.M.R., E.J.R., and J.P.C.; conceptualization: W.T.H., W.F., M.F., A.T.E., J.W.T., E.J.R., and J.P.C.; methodology: W.T.H., W.F., M.F., H.E.D., J.W.T., E.J.R., and J.P.C.; software: W.T.H., W.F., M.F., H.E.D., A.T.E., J.W.T., E.J.R., and J.P.C.; validation: W.T.H., W.F., M.F., A.T.E., J.W.T., E.J.R., and J.P.C.; formal analysis: W.T.H., W.F., M.F., J.W.T., E.J.R., and J.P.C.; investigation: W.T.H., W.F., M.F., S.M.R., D.W., A.M.H., and E.J.R.; resources: W.T.H., E.J.R., and J.P.C.; data curation: W.T.H., W.F., D.W., and E.J.R.; writing (original draft): W.T.H., W.F., and J.P.C.; writing (review and editing): W.T.H., W.F., M.F., A.S.A., H.E.D., S.M.R., D.W., A.M.H., A.T.E., J.W.T., E.J.R., and J.P.C.; visualization: W.T.H., W.F., M.F., A.T.E., J.W.T., E.J.R., and J.P.C.; supervision: J.W.T., E.J.R., and J.P.C.; administration: D.W., and J.P.C.; funding acquisition: W.T.H., W.F., M.F., H.E.D., A.T.E., J.W.T., E.J.R., and J.P.C.

## Supplemental Information

Supplement.pdf with Figures S1–5, Supplemental Table 1–2, and Supplemental Methods.

