## Supplement for "Unconstrained naturalistic human brain imaging and decoding with a fully wearable high- density optical system"

### Supplemental Information

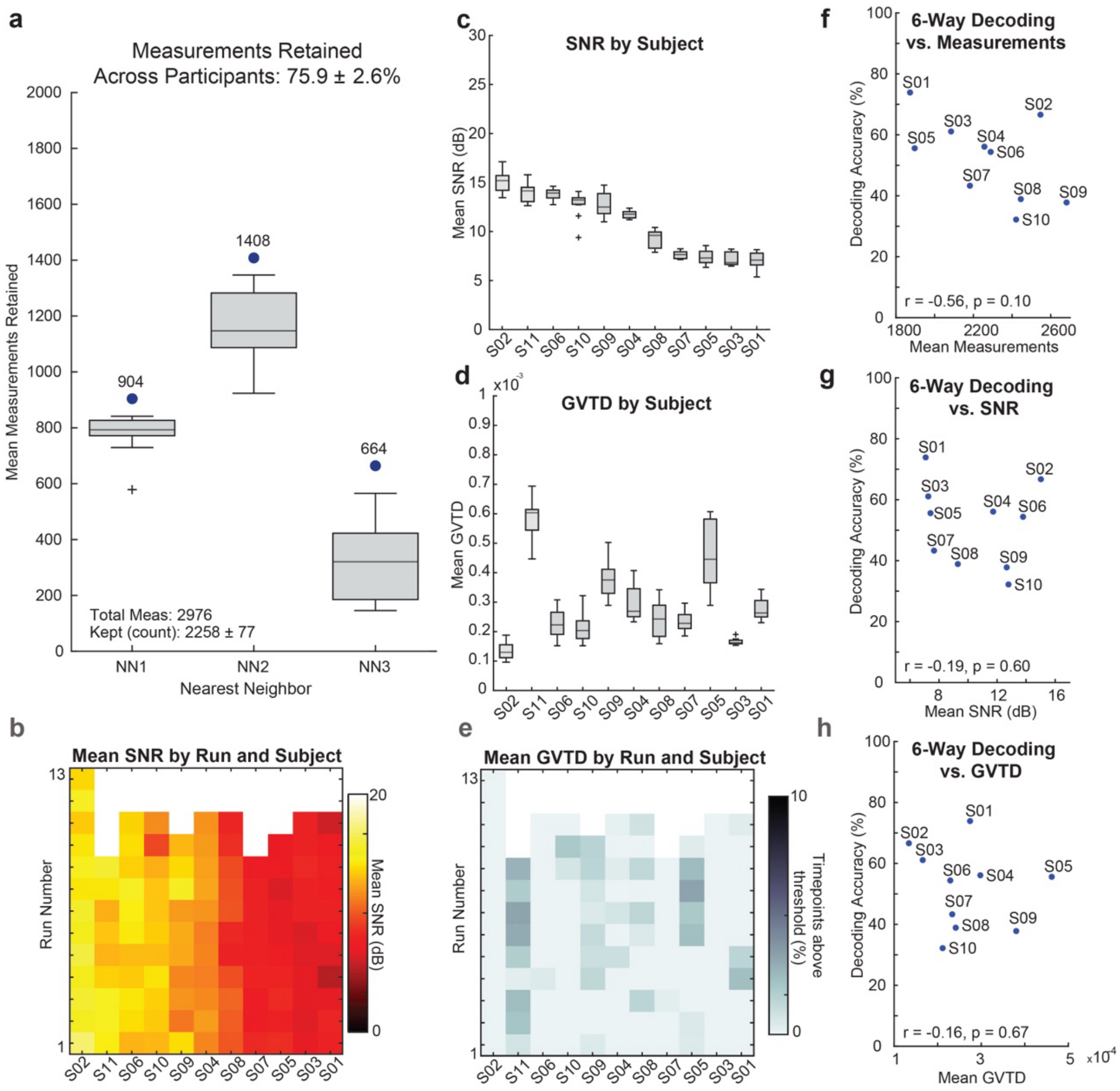


**Figure S1 | Data quality and motion metrics across participants and runs. (a)** Measurements retained across both wavelengths after quality filtering using a maximum source-detector separation of 40 mm and bad-channel exclusion methods (see **Methods** for details). Of 2,976 total measurements, an average of 2,258 ± 77 (mean ± SEM) measurements were retained across participants. **(b)** Heatmap of second nearest-neighbor (NN2) SNR across participants and runs. **(c)** Mean signal-to-noise ratio (SNR) of NN2 measurements at 850 nm across participants and runs shown as boxplots. Participants are ordered by decreasing mean SNR for visualization. **(d)** Global variance of the temporal derivatives (GVTD) motion metric across participants and runs. **(e)** Heatmap showing the percentage of time points retained above the GVTD threshold (1.5 × $10^{-3}$) across participants and runs. 6-Way localizer decoding performance vs **(f)** mean measurements retained, **(g)** mean pulse SNR (dB), **(h)** mean GVTD. **(a–e)** show data quality metrics for all participants (N=11), **(f–g)** show decoding performance vs data quality for participants with functional localizer analyses (N=10).

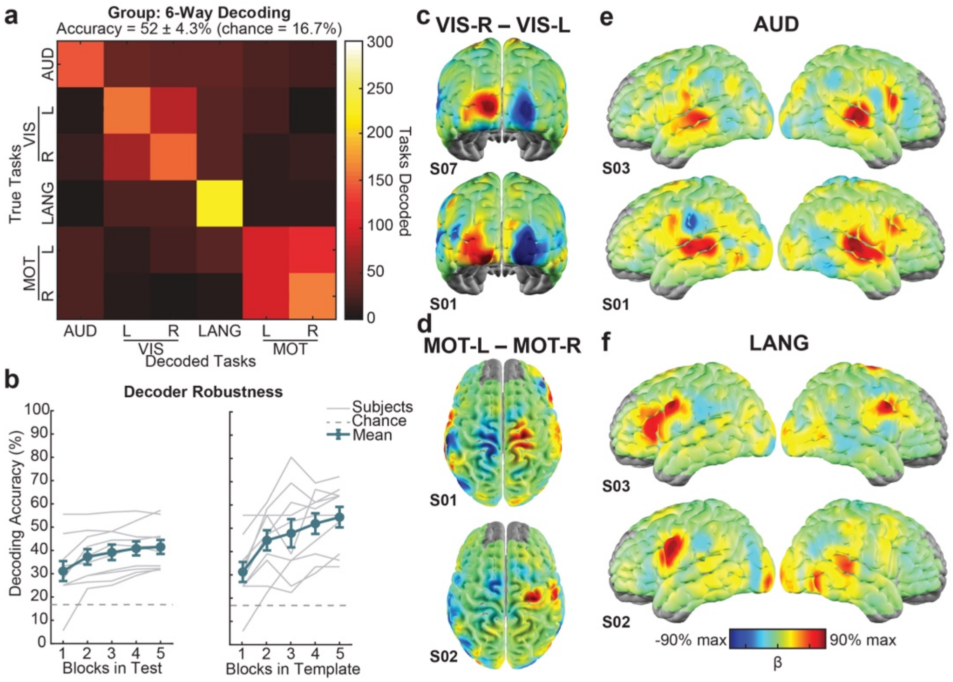


**Figure S2 | Group decoding robustness and single-participant functional localizer maps (HbO). (a)** Group-level decoding performance for the six-way functional localizer decoding task. **(b)** Decoder robustness analysis. Decoding performance was evaluated by independently varying the number of test blocks (left) and template blocks (right), while holding the complementary dataset constant at one block to isolate the effects of template and test data size. Decoding was performed in N=10. **(c–f)** GLM-derived activation maps from representative single participants for standard functional localizer tasks: **(c)** right vs left visual stimulation, **(d)** left vs right finger tapping, **(e)** auditory word presentation, and **(f)** a language task, demonstrating consistent spatial localization at the single-participant level.

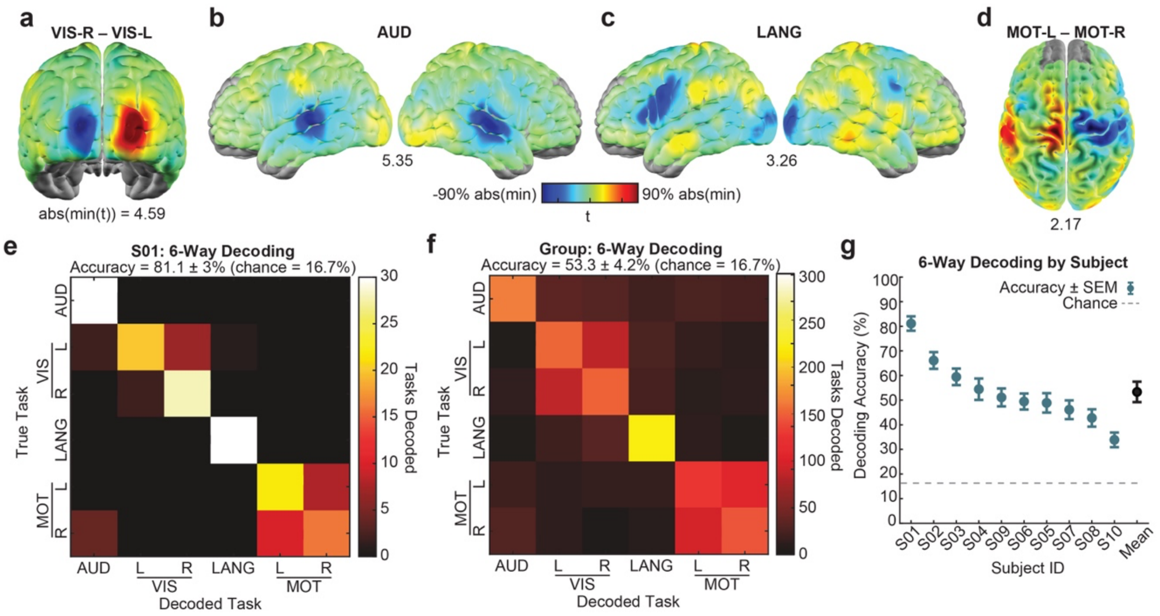


**Figure S3 | Group level localizer mapping and decoding performance using deoxyhemoglobin (HbR). (a–d)** Group fixed-effects t-statistic activation maps (N = 10 participants) derived from HbR responses for standard functional localizer tasks: **(a)** right vs left visual stimulation, **(b)** left vs right finger tapping, **(c)** auditory word presentation, and **(d)** language task. For visualization and comparison with HbO results, absolute t-values are shown. HbR t-values are comparable in magnitude to HbO (**Table 1**). **(e)** Representative single-participant decoding performance using HbR signals (mean accuracy 81.1 ± 3.0%), compared to HbO decoding (73.9 ± 3.0%; **Fig. 3d**) using the same template-based decoding framework. **(f)** Group-level decoding performance using HbR (53.3 ± 4.2%) compared to HbO (52.3 ± 4.3%; **Table 1**), demonstrating comparable decoding accuracy across chromophores. **(g)** Participant-level decoding accuracy using HbR signals. Performance varies across participants but remains above chance for all participants.

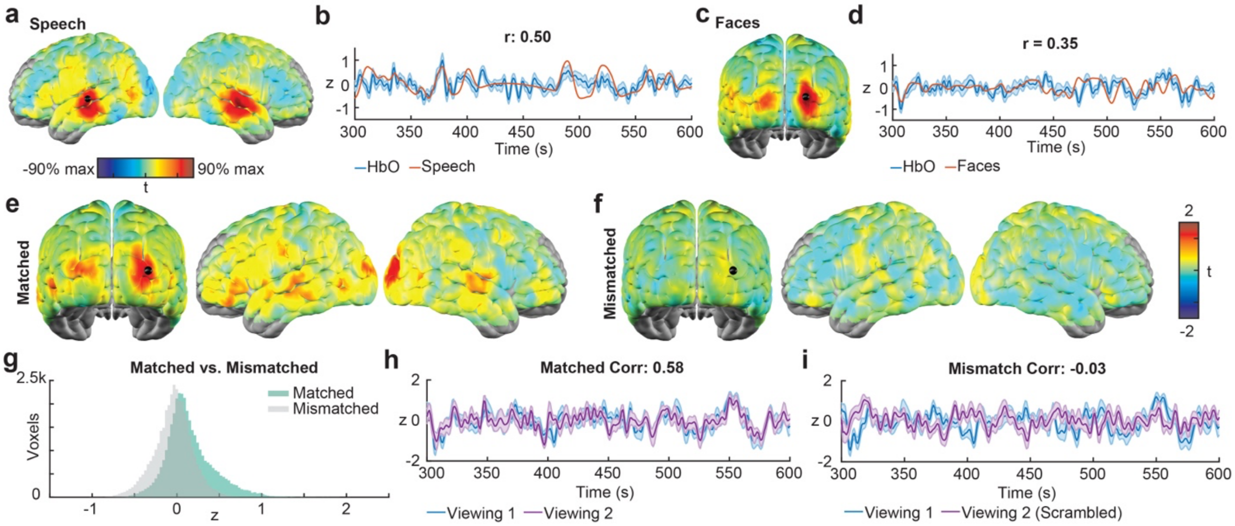


**Figure S4 | Naturalistic movie response synchronization and unfiltered feature analysis (HbO, N=10). (a–d)** Regressor-based movie feature analysis performed without additional temporal filtering. Spatial activation patterns and voxelwise correlations remain similar to those shown in **Fig. 5**, with slightly reduced correlation values. Representative correlations: speech (r = 0.50), faces (r = 0.35). **(e)** Voxelwise temporal correlation maps between first and second movie viewing runs (matched condition). Group t-statistic maps show synchronized responses across visual, temporal, and frontal cortical regions. **(f)** Voxelwise temporal correlation maps for mismatched movie segments. The second viewing was temporally reordered (halves swapped) to preserve autocorrelation structure while disrupting stimulus alignment. Group t-statistic maps show reduced or negative correlations distributed across cortex. **(g)** Histogram of voxelwise correlation values for matched and mismatched conditions derived from panels **(e)** and **(f)**, showing a rightward shift toward positive correlations for matched movie viewing. **(h–i)** Representative time courses from a voxel within the visual region for matched and mismatched comparisons. Matched responses show strong temporal alignment (r = 0.58), whereas mismatched responses show low correlation (r = −0.03).

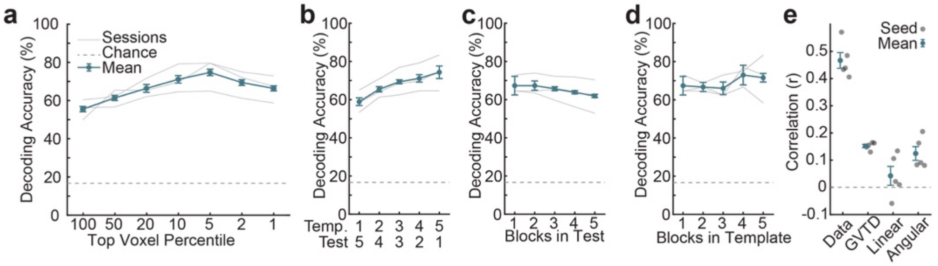


**Figure S5 | Naturalistic piano imaging robustness and paradigm structure. (a)** Varying thresholds for voxels included in decoding shows an increase in performance up to 5% voxels kept. **(b–d)** Piano decoder robustness with varying template/test blocks **(b)**, varying only test with one template block **(c)**, and varying only templates with one test block **(d)**. **(e)** Correlation of seed time traces and motion metrics, along with correlation between repeats of seed time trace data within sessions. Only correlation of seed time trace data shows high correlation, while correlation with motor metrics is low.

| System Specification | WHD-DOT | WHD-DOT Prototype | Fiber HD-DOT |
|---|---|---|---|
| **# Sources x Detectors** | 128 x 128 | 98 x 98 | 128 x 125 |
| **# Measurements (≤40mm)** | 2976 | 2320 | 3610 |
| **Nearest Neighbor Spacing (mm)** | 13 | 13 | 11 |
| **Wavelengths (nm)** | 735, 850 | 735, 850 | 685, 830 |
| **Detectivity (fW/[√Hz·mm$^2$])** | 11.2 | 10.5 | 10.6 |
| **Dynamic range (dB in 1 Hz)** | 151 | 155 | 156 |
| **Frame rate (Hz)** | 8 | 7.75 | 10.4 |

**Supplemental Table 1**. Comparison of system specifications for the wearable HD-DOT system developed in this study, with a first prototype of a wearable HD-DOT system developed in our lab as well as a representative fiber HD-DOT system. WHD-DOT Prototype: [1]. Fiber HD-DOT: [2].

| Subject | Hearing Words | Generate Verbs | Visual Checkerboard | Finger Tapping (Motor) | Movie Viewing | Piano Play |
|---|---|---|---|---|---|---|
| S01 | 4 | 2 | 2 | 1 | 2 | - |
| S02 | 4 | 4 | 2 | 1 | 2 | - |
| S03 | 4 | 2 | 2 | 1 | 2 | - |
| S04 | 4 | 2 | 2 | 1 | 2 | - |
| S05 | 4 | 1 | 2 | 1 | 2 | - |
| S06 | 4 | 2 | 2 | 1 | 2 | - |
| S07 | 3 | 1 | 2 | 1 | 2 | - |
| S08 | 4 | 2 | 2 | 1 | 2 | - |
| S09 | 4 | 1 | 2 | 1 | 2 | - |
| S10 | 4 | 2 | 2 | 1 | 2 | - |
| S11 | - | - | - | - | - | 9 |
| **DOT Total (N=11)** | 39 | 19 | 20 | 10 | 20 | 9 |
| **fMRI (Group, N=15)** | 64 | 48 | 39 | 39 | - | - |

**Supplemental Table 2**. This table contains the total number of runs in each of the 11 participants included in the data analysis. The WHD-DOT data were collected across two sessions in ten participants. Data for the piano study were collected across three imaging sessions in one participant. The MRI data are derived from previous studies performed in our lab.

### Supplemental Methods

Extended fMRI Preprocessing Details (from fMRIPrep)

The following fMRI preprocessing methods are derived from the fMRIPrep boilerplate text as recommended for citing fMRIPrep.

Preprocessing of B0 inhomogeneity mappings: Each subject had at least one field map per imaging session. A B0-nonuniformity map (or fieldmap) was estimated based on two (or more) echo-planar imaging (EPI) references with topup [3]; FSL 6.0.5.1:57b01774).

Anatomical data preprocessing: A total of 1 T1-weighted (T1w) images were found per subject within the input BIDS dataset. The T1-weighted (T1w) image was corrected for intensity non-uniformity (INU) with N4BiasFieldCorrection [4], distributed with ANTs 2.3.3 ([5], RRID:SCR_004757), and used as T1w-reference throughout the workflow. The T1w-reference was then skull-stripped with a Nipype implementation of the antsBrainExtraction.sh workflow (from ANTs), using OASIS30ANTs as target template. Brain tissue segmentation of cerebrospinal fluid (CSF), white-matter (WM) and gray-matter (GM) was performed on the brain-extracted T1w using fast (FSL 6.0.5.1:57b01774, RRID:SCR_002823, [6]. Brain surfaces were reconstructed using recon-all (FreeSurfer 7.2.0, RRID:SCR_001847, [7], and the brain mask estimated previously was refined with a custom variation of the method to reconcile ANTs-derived and FreeSurfer-derived segmentations of the cortical gray-matter of Mindboggle (RRID:SCR_002438, [8]. Volume-based spatial normalization to one standard space (MNI152NLin2009cAsym) was performed through nonlinear registration with antsRegistration (ANTs 2.3.3), using brain-extracted versions of both T1w reference and the T1w template. The following template was selected for spatial normalization: ICBM 152 Nonlinear Asymmetrical template version 2009c [[9], RRID:SCR_008796; TemplateFlow ID: MNI152NLin2009cAsym].

Functional data preprocessing: For each of the BOLD runs found per subject (across all tasks and sessions), the following preprocessing was performed. First, a reference volume and its skull-stripped version were generated by aligning and averaging 1 single-band references (SBRefs). Head-motion parameters with respect to the BOLD reference (transformation matrices, and six corresponding rotation and translation parameters) are estimated before any spatiotemporal filtering using mcflirt (FSL 6.0.5.1:57b01774, [10]). The estimated fieldmap was then aligned with rigid-registration to the target EPI (echo-planar imaging) reference run. The field coefficients were mapped on to the reference EPI using the transform. BOLD runs were slice-time corrected to 0.566s (0.5 of slice acquisition range 0s-1.13s) using 3dTshift from AFNI [11], RRID:SCR_005927). The BOLD reference was then co-registered to the T1w reference using bbregister (FreeSurfer) which implements boundary-based registration [12]. Co-registration was configured with six degrees of freedom. First, a reference volume and its skull-stripped version were generated using a custom methodology of fMRIPrep. Several confounding time-series were calculated based on the preprocessed BOLD: framewise displacement (FD), DVARS and three region-wise global signals. FD was computed using two formulations following Power (absolute sum of relative motions, [13]) and Jenkinson (relative root mean square displacement between affines, [10]). FD and DVARS are calculated for each functional run, both using their implementations in Nipype (following the definitions by [13]. The three global signals are extracted within the CSF, the WM, and the whole-brain masks. Additionally, a set of physiological regressors were extracted to allow for component-based noise correction (CompCor, [14]). Principal components are estimated after high-pass filtering the preprocessed BOLD time-series (using a discrete cosine filter with 128s cut-off) for the two CompCor variants: temporal (tCompCor) and anatomical (aCompCor). tCompCor components are then calculated from the top 2% variable voxels within the brain mask. For aCompCor, three probabilistic masks (CSF, WM and combined CSF+WM) are generated in anatomical space. The implementation differs from that of Behzadi et al. in that instead of eroding the masks by 2 pixels on BOLD space, a mask of pixels that likely contain a volume fraction of GM is subtracted from the aCompCor masks. This mask is obtained by dilating a GM mask extracted from the FreeSurfer's aseg segmentation, and it ensures components are not extracted from voxels containing a minimal fraction of GM. Finally, these masks are resampled into BOLD space and binarized by thresholding at 0.99 (as in the original implementation). Components are also calculated separately within the WM and CSF masks. For each CompCor decomposition, the k components with the largest singular values are retained, such that the retained components' time series are sufficient to explain 50 percent of variance across the nuisance mask (CSF, WM, combined, or temporal). The remaining components are dropped from consideration. The head-motion estimates calculated in the correction step were also placed within the corresponding confounds file. The confound time series derived from head motion estimates and global signals were expanded with the inclusion of temporal derivatives and quadratic terms for each [15]. Frames that exceeded a threshold of 0.5 mm FD or 1.5 standardized DVARS were annotated as motion outliers. Additional nuisance timeseries are calculated by means of principal components analysis of the signal found within a thin band (crown) of voxels around the edge of the brain, as proposed by [16]. The BOLD time-series were resampled into standard space, generating a preprocessed BOLD run in MNI152NLin2009cAsym space. First, a reference volume and its skull-stripped version were generated using a custom methodology of fMRIPrep. All resamplings can be performed with a single interpolation step by composing all the pertinent transformations (i.e. head-motion transform matrices, susceptibility distortion correction when available, and co-registrations to anatomical and output spaces). Gridded (volumetric) resamplings were performed using antsApplyTransforms (ANTs), configured with Lanczos interpolation to minimize the smoothing effects of other kernels [17]. Non-gridded (surface) resamplings were performed using mri_vol2surf (FreeSurfer).
